# A Multi-Tetracycline Responsive Induction System for Gene Expression in *Bacillus subtilis*

**DOI:** 10.1101/2025.09.17.676849

**Authors:** Keira Reich-Veillette, Elizabeth A. Libby

**Affiliations:** Department of Bioengineering, Northeastern University, Boston MA 02115 USA

**Keywords:** *Bacillus subtilis*, tetracycline-inducible system, anhydrotetracycline, chlortetracycline, doxycycline

## Abstract

Well-characterized induction systems that offer tightly regulated, graded, and homogeneous control of gene expression are essential tools for basic and applied microbiology. Tetracycline-based induction systems are widely used non-toxic expression systems in bacteria, however the GRAS and industrially relevant model organism *Bacillus subtilis* currently lacks a tightly regulated, sensitive system that is robust to media and growth conditions. Here we adapted features of existing tetracycline induction systems to the specific requirements of *B. subtilis* by engineering a codon-optimized TetR repressor, enhanced promoter and operator architecture, and a modular shuttle vector enabling stable, single-copy chromosomal integration. The resulting system achieves ∼1000-fold dynamic range with minimal leakiness, using subinhibitory concentrations of anhydrotetracycline, chlortetracycline, and doxycycline as inducers in rich and minimal media as well as in colonies. We further quantify the titration parameters and non-inhibitory ranges for each inducer and growth condition, providing a well-characterized and versatile tool for inducible gene expression in *B. subtilis*.

**Importance:** *Bacillus subtilis* is an important bacterial model organism with applications in biotechnology and industrial microbiology. However, tools for tightly controlled gene regulation that are robust to growth conditions remain limited. We developed a tetracycline-inducible system tailored to *B. subtilis,* providing strong, tunable expression with minimal leaky expression across diverse growth conditions. Due to its enhanced sensitivity, tight regulation, and homogeneous induction of the bacterial population, it enables precision titration of gene expression for a variety of downstream applications such as physiology studies, pathway optimization, biotechnology, and colony-based screening.

## INTRODUCTION

Induction systems for tightly and robustly regulating bacterial gene expression are essential tools in microbiology and biotechnology research. Established systems with diffusible, non-inhibitory, and non-metabolized inducers, such as IPTG for the *lac* operon, are particularly valuable for their predictable expression. To date, many inducible systems and regulatory elements have been developed for bacterial systems, particularly the model organisms. The model organism *Escherichia coli* has many well-characterized induction systems, however, there are currently more limited options for *Bacillus subtilis* (*1, 2*).

*B. subtilis* is a naturally competent and a generally recognized as safe (GRAS), industrially important model organism with strong protein secretion capabilities, relative ease of genetic manipulation, and exhibits prototypical developmental processes such as sporulation and biofilm formation (*2–7*). An increasing number of tools have been developed for *B. subtilis* in recent years, including modular shuttle vectors (*8*), native and synthetic promoter libraries (*9, 10*), and RBS toolboxes (*11, 12*). However, a major gap remains in the development of well-characterized inducible control systems that use non-inhibitory and non-metabolized inducers. Currently, the most widely used inducers include IPTG (*13–15*), xylose (*15, 16*), mannose (*17, 18*), maltose (*19*), cumate (*20*), theophylline (*21*), and bacitracin (*22, 23*). However, the large number of metabolizable or inhibitory inducers pose challenges in applications requiring multiple, media-independent systems, given the close interplay between metabolism and *B. subtilis* physiology (*24, 25*).

In *E. coli* and other organisms, Tn10 *tet* operon-based tetracycline-induction systems provide tight and non-metabolized inducer-based regulation at subinhibitory concentrations (*26–31*). In its native context, the *tet* operon confers resistance in many eubacteria to tetracycline antibiotics, which inhibit protein translation (*32*). The operon encodes the Tet repressor (TetR) which, when bound to tetracycline, relieves repression of downstream resistance genes, e.g., tetracycline antiporter gene *tetA* (*32–37*). Synthetic variations of this operon are desirable for their inherent high sensitivity and tight repression, attributes arising from the combined necessity to provide resistance at non-lethal tetracycline concentrations while preventing toxic levels of TetA expression (*37–39*).

Despite these advantages, no tetracycline-inducible system has been widely adopted for *B. subtilis*. Although *B. subtilis* harbors its own tetracycline resistance system (*40*), it lacks the canonical repressor-operator architecture found in *E. coli*, requiring synthetic designs to adapt *E. coli*-derived components. However, adapting biological parts with quantitative precision to a different species is challenging due to fundamental differences in cellular permeability to the inducer, regulatory architectures, codon usage, and transcriptional machinery. These factors all affect system performance and likely contribute to the limited progress in *B. subtilis,* which is more sensitive to tetracyclines, and is also particularly sensitive to promoter element composition and ribosomal binding site (RBS) positioning (*41, 42*). The first tetracycline-inducible system for *B. subtilis* was developed over 30 years ago (*43, 44*), and although the biological parts have since been adapted for other gram-positive bacteria (*45–48*), efforts to improve this system to operate in *B. subtilis* have remained sparse. Early studies primarily focused on expanding the attainable regulatory window, resulting in a dynamic range of ∼300-fold to anhydrotetracycline (aTc) (*49, 50*).

More recent studies characterizing dose-response behavior of tetracycline-inducible systems in *B. subtilis* have highlighted both the potential and the limitations of the current state of the art (*11, 51*). Some system variants displayed titratable expression, yet others still largely measured the commonly used aTc-mediated induction as largely all-or-none, limiting fine-tuned control. These systems also exhibited practical constraints including short induction windows, population heterogeneity, and qualitatively inconsistent dose-responses to inducer across media types. Additionally, to our knowledge, there are no reports of successful use of these systems to induce gene expression in colonies grown on agar plates, which would facilitate important applications such as library screens and selections.

Many of the challenges experienced with the existing systems could stem from the biological parts requiring further *B. subtilis*-specific adaptation. Close examination of the promoter architecture suggested it could be an important variable for achieving both tight repression and high dynamic range. Many existing designs favor a single operator site, as the dual operators reduced system leakiness in the absence of inducer but often at the cost of maximal potential expression (*43, 49, 51*). These observations suggested that carefully tailoring the induction circuit could offer a large dynamic range, high sensitivity, and tunable graded control, while exploring colony-level induction addresses an important gap, ultimately enabling a broadly useful system for *B. subtilis*.

We developed a titratable tetracycline-inducible system tailored to *B. subtilis* that can achieve a ∼1000-fold dynamic range in liquid culture. The system employs the TetR(B) repressor and a *P_tet_* promoter on a modular shuttle vector designed for single-copy chromosomal integration. This system exhibits measurable response to aTc induction at a concentration two orders of magnitude lower than those reported in previous work, limiting inducer toxicity. The high sensitivity enables system compatibility with two additional tetracyclines: chlortetracycline (cTc) and doxycycline (dox), inducers not typically implemented in bacterial systems due to their higher antibiotic activity. The different affinities of these tetracyclines for TetR enable unique induction profiles (*52, 53*), providing fine-tuned titratability. We present a quantitative characterization of this system with these three tetracycline inducers across diverse growth conditions, including rich and minimal liquid media and in colonies, expanding the toolkit of well-characterized inducible systems in *B. subtilis*.

## RESULTS

### Design of a tightly regulated tetracycline induction circuit for integration into *B. subtilis* chromosome

We sought to create a tightly regulated tetracycline induction circuit for use in *B. subtilis* by adapting and editing existing Tn10 regulatory elements. The *tetR(B)* gene was codon-optimized for *B. subtilis* and placed in front of a constitutive penicillinase promoter (P*_pcn_*) (*13*) (Fig. 1A). Two tet operator sites (*tetO_1_* and *tetO_2_*) were combined with promoter elements for σ^A^, permitting expression in vegetative growth, to create an inducible *P_tet_* with tight regulation. For quantitative characterization of this system, we used this promoter to regulate the expression of *mcherry*. This system was integrated into a shuttle vector for double crossover recombination into the *B. subtilis* chromosome at the *amyE* locus with a spectinomycin resistance marker.

**Figure 1.**
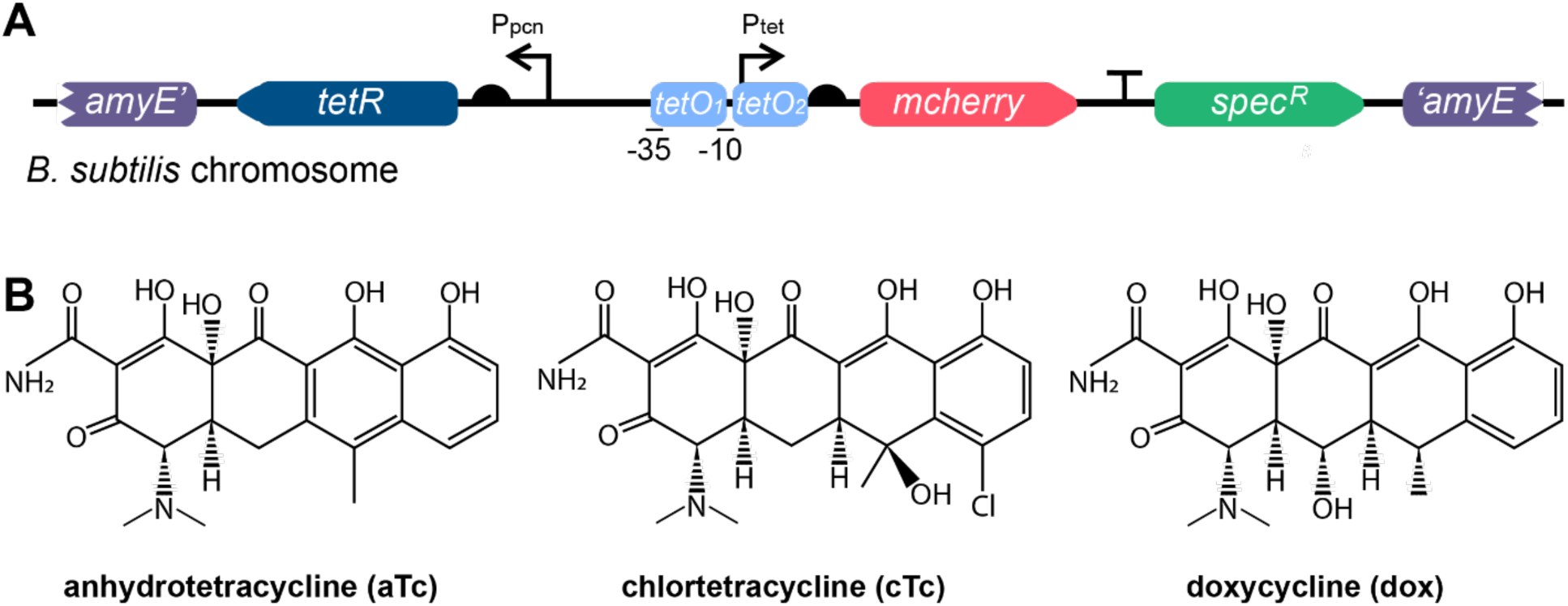
Design of tetracycline inducible system in *B. subtilis*. (A) A codon-optimized *tetR* is expressed under a constitutive promoter (P*_pcn_*) with a strong ribosome binding site (RBS) with a 9-bp spacer. The reporter gene *mcherry* is expressed from a regulatory element consisting of a P*_tet_* promoter with two operator sites (*tetO_1_* and *tetO_2_*) and a -10/-35 of *TACAAT* / *TTGACA*. The induction system is cloned into a shuttle vector for single copy integration into the *B. subtilis* chromosome at the *amyE* locus. (B) Chemical structures of the tetracycline family antibiotics anhydrotetracycline (aTc), chlortetracycline (cTc) and doxycycline (dox).

In this design, in the absence of tetracycline, TetR binds the *tetO* sites repressing *P_tet_* and preventing expression of *mcherry*. Once added, tetracycline binds to TetR, resulting in loss of repression, inducing *mcherry* expression and subsequent fluorescence. Different tetracycline family antibiotics have varied affinities for TetR, resulting in different dose-response induction curves. Therefore, we sought to characterize our improved circuit design with three tetracycline family antibiotics; anhydrotetracycline, chlortetracycline and doxycycline (Fig. 1B), that are commonly used in a variety of other organisms as inducers and have superior induction compared to tetracycline (*52, 53*).

### Characterization of the tetracycline inducible system in minimal media

We first sought to measure the performance of our designed tetracycline induction system in minimal glucose media with respect to its leakiness, sensitivity, uniformity, and stability throughout growth. This was done for each of the tetracycline family antibiotics: aTc, cTc, and dox as detailed individually below. For all of these inducers, we measured mCherry fluorescence throughout growth in varying concentrations of each tetracycline (Fig. S1). Dose-response curves were generated using the fluorescent signal at the ∼ start of log phase for each respective concentration (Fig. 2A-C). Importantly, we did not observe any consistently detectable signal over cellular autofluorescence for the system in the absence of tetracycline under these conditions. This demonstrates stable and tight regulation of this system in minimal glucose media but prevents the calculation of a true fold-change induction under these conditions.

**Figure 2.**
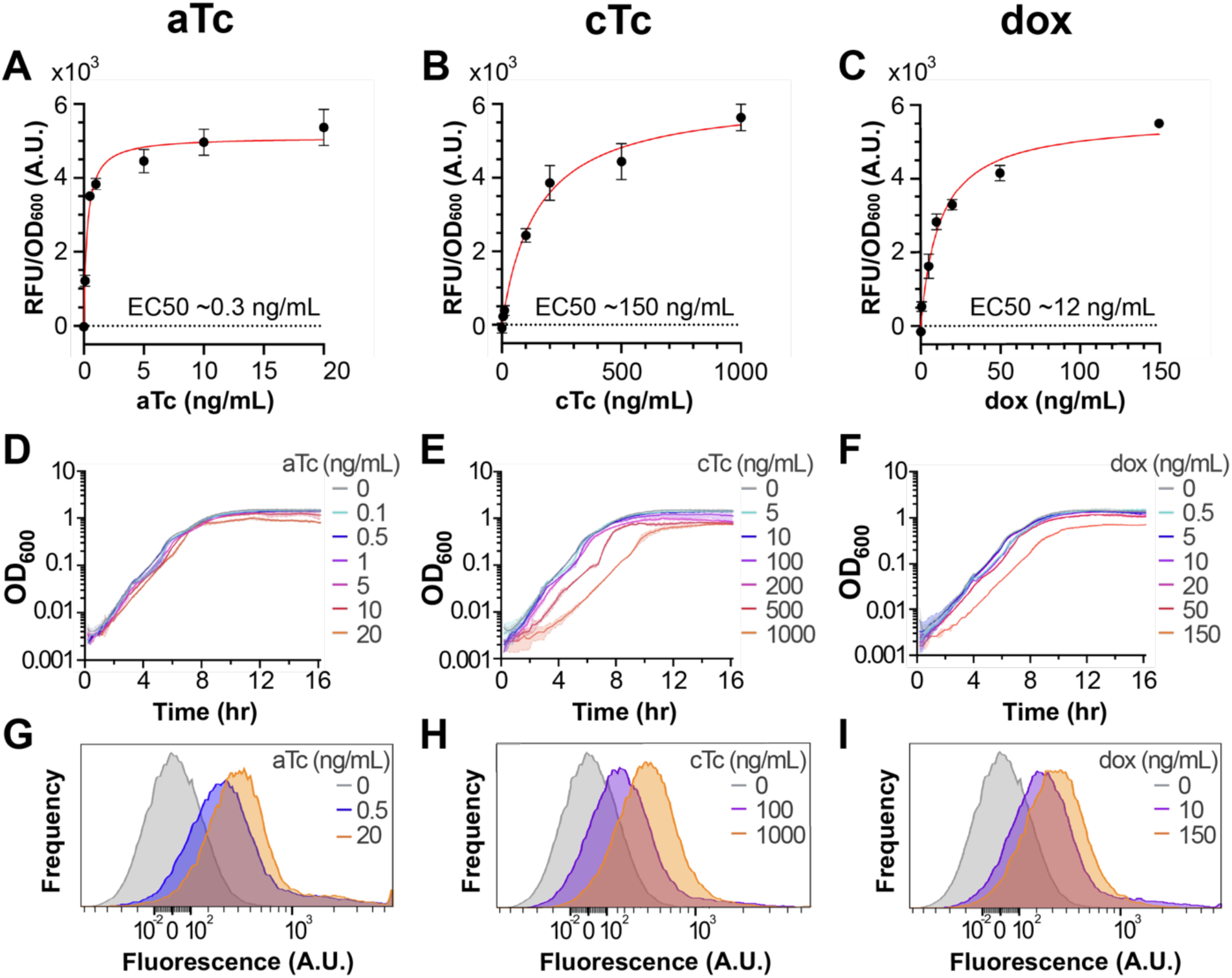
Tetracycline induction in minimal glucose media. A-C: Dose-response induction using (A) aTc, (B) cTc and (C) dox at ∼ mid-log (OD_600_ ∼ 0.3) in minimal glucose media. Red: Nonlinear regression fits with approximate EC50 values as indicated. Dots and bars indicate the mean and SEM respectively of three biologically independent experiments. D-F: Corresponding growth curves with (D) aTc, (E) cTc and (F) dox from representative experiments. Shading represents standard deviation (SD) across triplicates within a single experiment. G-I: Measurements of mCherry fluorescence distributions by flow cytometry induction from at least 40,000 events for (G) aTc, (H) cTc and (I) dox. Concentrations at ∼ each respective tetracycline’s EC50 and its maximal tested concentration are shown, with the 0 ng/mL condition shown for reference.

For each tetracycline, growth curves were measured (Fig. 2D-F) and the effects of each inducer concentration on bacterial growth rate were quantified by comparing doubling times to that without inducer (Table 1). To assess the homogeneity of the fluorescence distributions in single-cells, we used flow cytometry (Fig. 2G-I). Similarly to the bulk culture measurements, we did not observe any significant leaky expression in the absence of tetracycline.

**Table 1.**
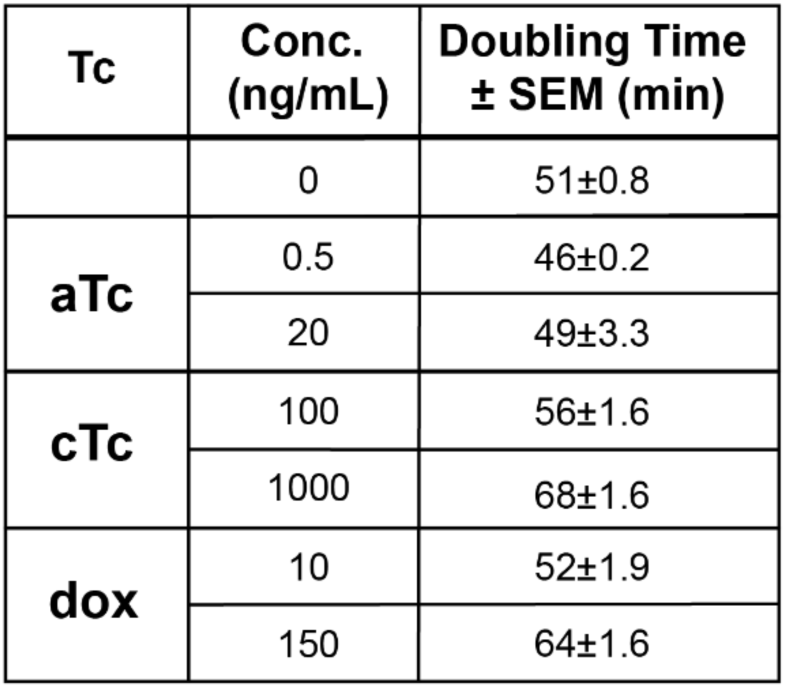
Doubling times for cultures grown in minimal glucose media induced with tetracyclines. Doubling times in log phase (up to OD_600_ ∼ 0.15) for cultures grown in minimal glucose media and induced with tetracyclines aTc, cTc, and dox. Concentrations at each respective tetracycline’s ∼ EC50 and its maximal tested concentration are shown.

#### Anhydrotetracycline

Anhydrotetracycline, or aTc, is one of the most widely used inducers for tetracycline-responsive systems. Therefore, we titrated aTc from 0 – 20 ng/mL, a non-toxic range for *B. subtilis*. We observed a maximum mCherry signal >5100 RFU/OD_600_, using 10-20 ng/mL of aTc, compared with ∼ 0 without inducer (Fig. 2A). We measured induction using aTc concentrations as low as 0.1 ng/mL with half-max (EC50) at ∼0.3 ng/mL. The growth curves remain similar across all tested concentrations (Fig. 2D), and calculated doubling times in log phase are not significantly affected at 20 ng/mL, the highest concentration (Table 1). However, earlier stationary phase lysis is observed using 20 ng/mL, potentially indicating a mild delayed toxicity of unknown origin (Fig. S1). Single-cell fluorescence distributions measured by flow cytometry show unimodal expression at representative inducer concentrations (Fig. 2G). This unimodality was also observed at the lower doses, demonstrating that the system maintains predictable, population-wide inducibility even at low concentrations. Geometric mean fluorescence increased from 13.7 ± 0.4 in the uninduced control to 330 ± 0.7 at 0.5 ng/mL and 431 ± 4.2 at 20 ng/mL, with error reported as the standard deviation of separately induced replicates. Limited autofluorescence was observed in the non-fluorescent control strain treated with aTc at identical dosages (Fig. S2).

#### Chlortetracycline

Chlortetracycline, or cTc, has a lower affinity for TetR than aTc, suggesting it likely requires a higher amount of antibiotic to induce P*_tet_*. Therefore, we used a larger range of inducer concentrations to measure its full dynamic range: 0-1000 ng/mL. Half-max induction (EC50) occurred at ∼150 ng/mL (Fig. 2B). Although the dose-response curve predicts a maximal mCherry signal > 6200 RFU/OD_600_, this level is not experimentally attainable due to high toxicity; ∼5600 RFU/OD_600_ at 1000 ng/mL was the highest signal achieved. This toxicity is reflected in significant perturbations to growth, including a long lag phase (Fig. 2E) and increases in doubling times up to ∼30% at 1000 ng/mL cTc (Table 1). However, single-cell fluorescence distributions still show unimodal expression, including at lower inducer concentrations tested (Fig. 2H). Geometric mean fluorescence increased from 13.7 ± 0.4 in the uninduced control to 195 ± 8.5 at 100 ng/mL and 346 ± 4.2 at 1000 ng/mL, with error as the standard deviation of separately induced replicates. Like aTc, little to no observable autofluorescence is measured in the non-fluorescent control treated with cTc at matched inducer concentrations (Fig. S2).

#### Doxycycline

Doxycycline is not as commonly used to induce P*_tet_* in bacterial systems as it is in eukaryotic systems. Therefore, we empirically tested the response of our induction system to a range of dox concentrations. We found ∼0-150 ng/mL allowed us to observe a full range of induction, suggesting it has intermediate potency compared to aTc and cTc under these growth conditions. The dose-response estimates a half-max signal (EC50) at ∼12 ng/mL and maximal induction of >5600 RFU/OD_600_ at ∼150 ng/mL (Fig. 2C). However, growth is impacted by this concentration, including a longer lag (Fig. 2F) and a ∼25% increase in doubling time (Table 1). Toxicity appears to be much lower in the rest of the tested concentrations, including 50 ng/mL: ∼10% increase in doubling time while achieving ∼75% of maximal induction. This demonstrates that intermediate concentrations of dox can produce an easily titratable signal to nearly full induction. Similarly to aTc and cTc, dox resulted in unimodal induction of the population as measured by flow cytometry at all tested concentrations (Fig. 2I). Geometric mean fluorescence increased from 13.7 ± 0.4 in the uninduced control to 262 ± 8.5 at 10 ng/mL and 321 ± 3.5 at 150 ng/mL, with error as the standard deviation of separately induced replicates. Limited autofluorescence was observed in the non-fluorescent control strain treated with dox at matched inducer concentrations (Fig. S2).

### Measurement of tetracycline system induction in colonies

Having established induction measurements for our system in liquid culture, we next sought to evaluate its performance in colonies grown on solid minimal glucose media (Materials and Methods). To do so, small amounts of *B. subtilis* cultures were spotted on minimal glucose agar plates containing the tetracyclines, and the resulting colonies were imaged for mCherry fluorescence (Fig. 3A). For each tetracycline, we used a concentration ∼EC50 and maximum inducer established in liquid above. This is with the exception of cTc, as toxicity at 1000 ng/mL inhibited colony growth, so 500 ng/mL cTc was used instead. A nonfluorescent wild-type strain (*Bs*168) colony was grown side-by-side with the strain containing the tetracycline inducible system to measure the contribution of each tetracycline to colony autofluorescence. Additionally, an agar plate without added inducer was included to measure the leakiness of this *P_tet_* system in colonies.

**Figure 3.**
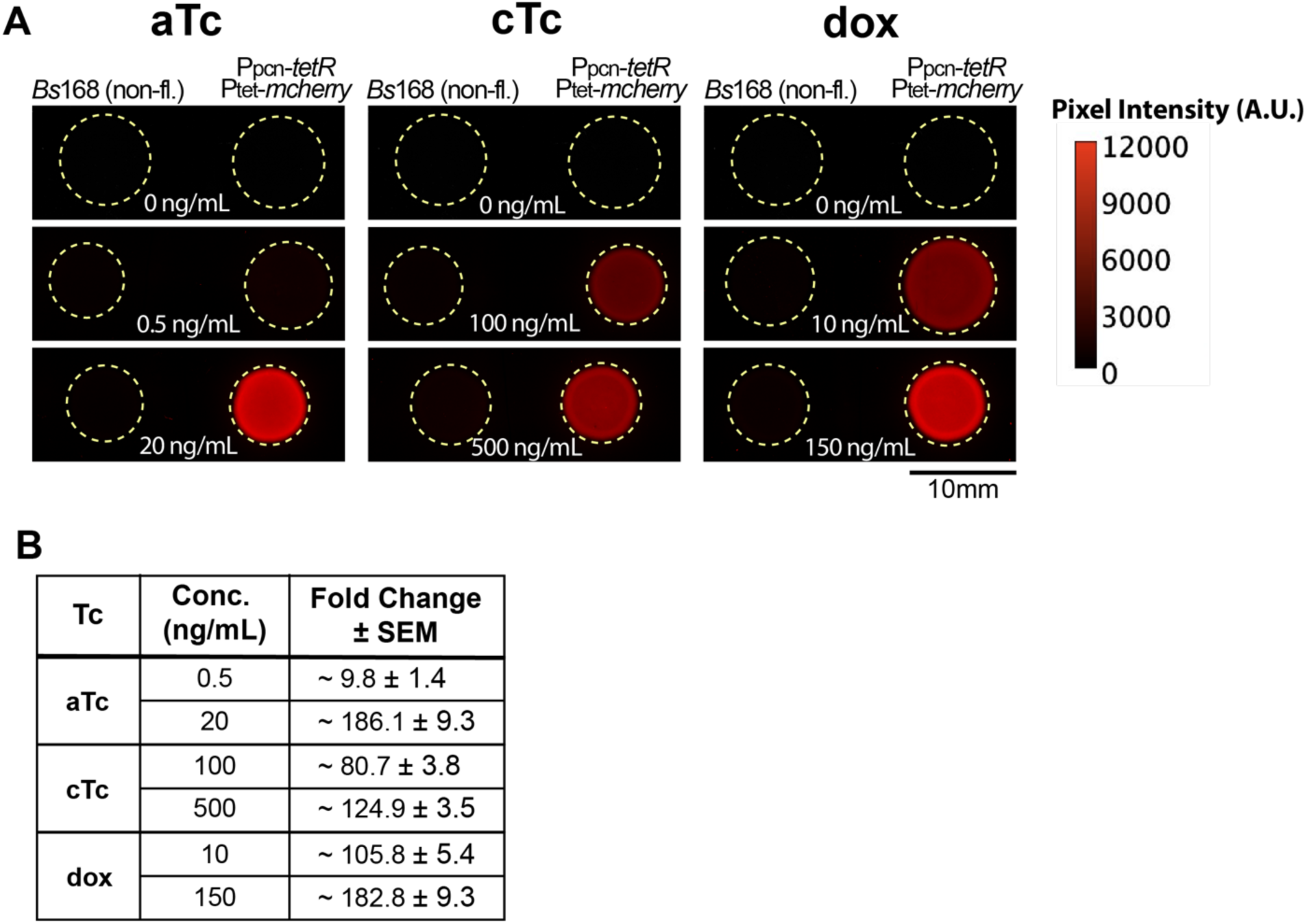
Tetracycline induction on colonies grown on solid media. (A) Representative mCherry fluorescence of *B. subtilis* colonies grown on minimal glucose agar plates supplemented aTc, cTc and dox. Concentrations at each tetracycline’s ∼ EC50 and its maximal tested concentration are shown, with the 0 ng/mL condition shown for reference across all tetracyclines. Dashed yellow circles indicate colony boundaries (Fig. S3, Brightfield). (B) Fold change in mean fluorescence calculated for each tetracycline from at least 3 biologically independent experiments.

In the absence of tetracycline, we observed little detectable colony fluorescence (51.4 ± 7.7 RFU), indicating ∼tight repression of mCherry in our system, but considerably more leaky expression than observed in liquid culture. Colony fluorescence, mCherry, increased with higher concentrations of each tetracycline inducer, demonstrating the ability to use this system at a macro scale. To quantify this response, we measured pixel intensity in the mCherry channel for each colony. From these values, the intensity fold change was calculated relative to colonies without inducer (Figure 3B). Maximum concentrations for aTc and dox resulted in fold changes of ∼200 fold, and the 500 ng/mL cTc resulted in ∼150-fold induction.

### Characterization of the tetracycline inducible system in rich media

With system induction characterized in minimal media, we extended our analysis to a commonly used rich media (CH) to confirm general induction parameters remain qualitatively similar. We repeated the above strategy and measured dose-response induction to each tetracycline. We found that the induction profiles were qualitatively similar to those observed in minimal glucose media (Figure 4). While overall observed maximal induction was reduced by up to ∼50% and half-max values shifted down by ∼50-80% depending on the tetracycline, the absence of leaky expression was maintained, demonstrating its utility across nutrient conditions.

**Figure 4.**
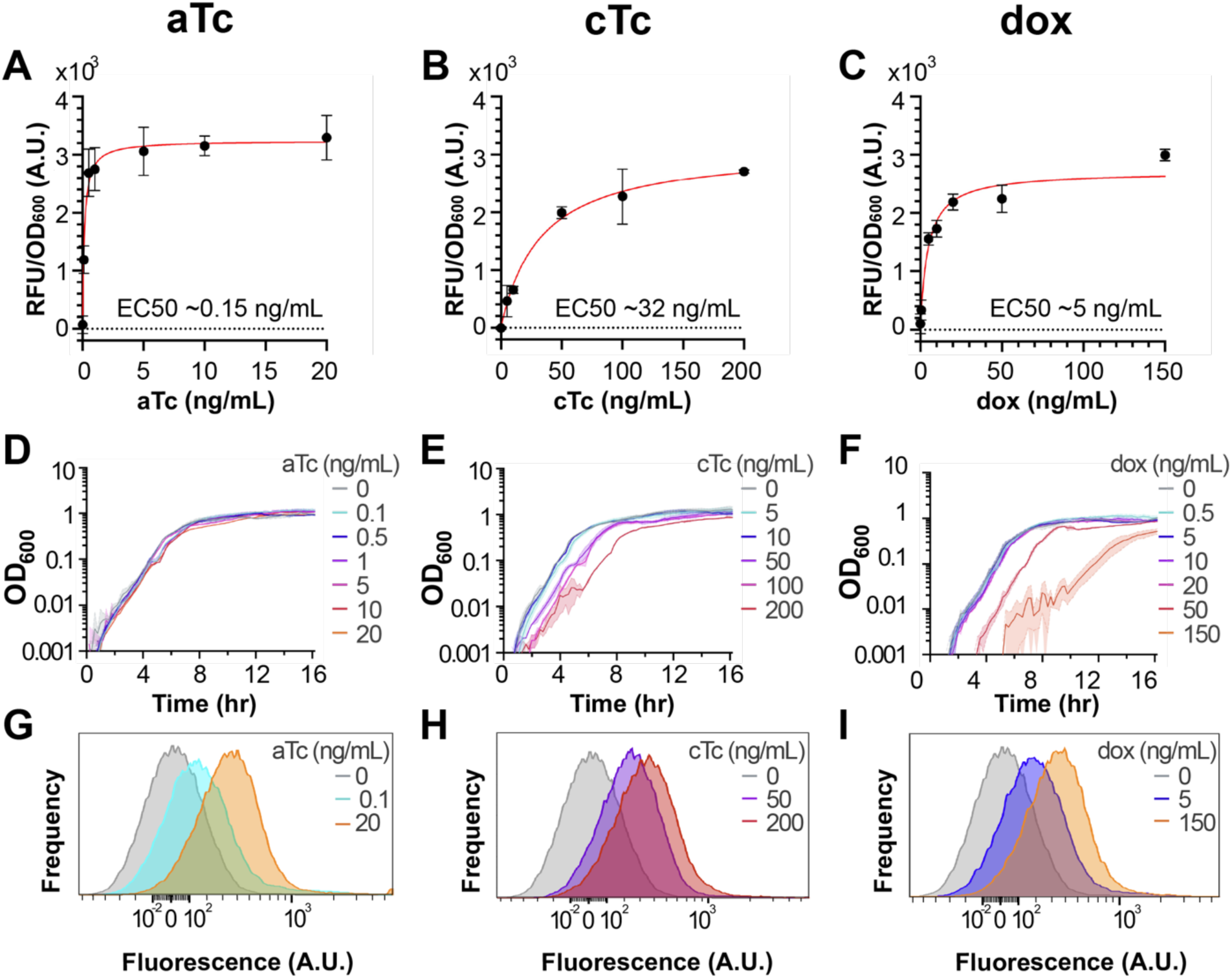
Tetracycline induction in rich media. A-C: Dose-response induction using (A) aTc, (B) cTc and (C) dox at ∼ mid-log (OD_600_ ∼ 0.3) in CH media. Red: Nonlinear regression fits with approximate EC50 values as indicated. Dots and bars indicate the mean and SEM respectively of three biologically independent experiments. D-F: Corresponding growth curves with (D) aTc, (E) cTc and (F) dox from representative experiments. Shading represents standard deviation (SD) across triplicates within a single experiment. G-I: Measurements of mCherry fluorescence distributions by flow cytometry induction from at least 40,000 events for (G) aTc, (H) cTc and (I) dox. Concentrations at ∼ each respective tetracycline’s EC50 and its maximal tested concentration are shown, with the 0 ng/mL condition shown for reference.

#### Anhydrotetracycline

To facilitate direct comparison of induction parameters to minimal media, matched aTc concentrations were used in rich media. A fit to the dose-response data shows a maximum induction of >3200 RFU/OD_600_ with half-max signal (EC50) at ∼ 0.15 ng/mL aTc (Fig. 4A). aTc showed a slight impact on growth rate and lag at its higher concentrations (Figure 4D), with doubling times (Table 2) increasing by ∼10% at 10 ng/mL. Single-cell fluorescence distributions measured by flow cytometry (Figure 4G) show unimodal expression at all concentrations, indicating population-wide induction, and limited autofluorescence was observed in the non- fluorescent control strain at identical dosages (Fig. S5). Geometric mean fluorescence increased from 26 ± 0.5 in the uninduced control to 166 ± 0 at 0.1 ng/mL and 349 ± 4.2 at 20 ng/mL, with error as the standard deviation of separately induced replicates.

**Table 2.**
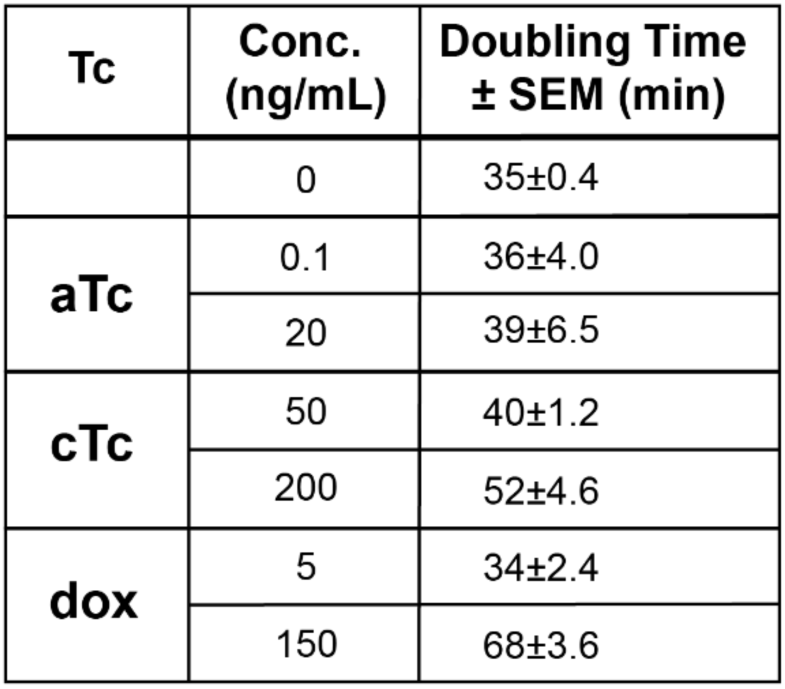
Doubling times for cultures in rich media with tetracyclines. Doubling times in log phase (OD600 ∼0.01-0.4) for cultures grown in CH media supplemented with tetracyclines aTc, cTc, and dox. Concentrations at each respective tetracycline’s ∼ EC50 and its maximal tested concentration are shown.

#### Chlortetracycline

Consistent with the generally observed increases in toxicity in rich media, we had to exclude the highest concentrations of cTc, 500 and 1000 ng/mL, due to lack of growth. Therefore, we included a data point at 50 ng/mL cTc to more accurately measure the dose-response behavior in the non- toxic range. We found that ∼200 ng/mL cTc yielded the highest induction without lethality at ∼2700 RFU/OD_600_ (Fig. 4B). However, around the EC50, 50 ng/mL, we still observe measurable changes in growth (Fig. 4E). At this concentration, the doubling time increases by ∼15% and increases further to ∼50% at 200 ng/mL (Table 2). However, single-cell fluorescence distributions measured by flow cytometry still appear unimodal at all tested concentrations (Fig. 4H), and limited autofluorescence was observed in the non-fluorescent control strain at matched inducer concentrations (Fig. S5). Geometric mean fluorescence increased from 26 ± 0.5 in the uninduced control to 240 ± 2.8 at 50 ng/mL and 340 ± 22.6 at 200 ng/mL, with error as the standard deviation of separately induced replicates.

#### Doxycycline

While doxycycline was significantly more toxic to *B. subtilis* in rich media than in minimal, in contrast to cTc, the full range of inducer concentrations used in minimal media could be tested. A fit to the dose-response data shows maximum induction of ∼3000 RFU/OD_600_ with a EC50 of ∼5 ng/mL (Fig. 4C). However, we found that *B. subtilis* growth is significantly impacted at concentrations nearing full induction: by 50 ng/mL dox, we observe a significant lag phase (Fig. 4F), and by the maximum tested concentration of 150 ng/mL, there is a >90% increase in doubling time (Table 2). As with the other inducers, despite the toxicity, single-cell fluorescence distributions measured by flow cytometry show unimodal expression (Fig. 4I) with limited autofluorescence contributions from doxycycline at matched inducer concentrations (Fig. S5). Geometric mean fluorescence increased from 26 ± 0.5 in the uninduced control to 194 ± 4.2 at 5 ng/mL and 340 ± 4.9 at 150 ng/mL, with error as the standard deviation of separately induced replicates.

In both minimal and rich media, this tetracycline-inducible system for *B. subtilis* demonstrated tight regulation in the absence of inducer, with fluorescence measurements without inducer consistently approaching the detection threshold of cellular autofluorescence. Additionally, this system exhibits unimodal induction across all concentrations of the three tetracycline inducers tested. Induction increases the fluorescence from ∼ 0 to ∼ 3000-6000 RFU/OD6_00_, in liquid media, providing strong regulation and a large dynamic range. When compared with minimal media, in rich media, overall maximum signals were reduced by ∼35-50% and EC50 values shifted down, indicating higher sensitivity to the inducer but lower maximal output. Across all inducers, toxicity increased in rich media compared to minimal glucose. These results indicate that the system maintains low leakiness and consistent tunable induction properties across nutrient conditions, however media composition influences both inducer potency and toxicity profiles. Interestingly, the fold changes measured by flow cytometry were generally lower than those observed with the plate reader, which may be due to the more precisely matched excitation and emission filters available on the plate reader. Additionally, while plate reader measurements suggested reduced inducibility in enriched media compared to minimal media, this difference was not apparent in flow cytometry, where induction was highly comparable across media types. Together, these discrepancies likely reflect differences in instrument sensitivity, detection range, and potentially the way population-averaged signals (plate reader) versus single-cell distributions (flow cytometry) are captured.

## DISCUSSION

The tetracycline induction system described here tightly regulates gene expression in *B. subtilis*, achieving ∼1000-fold dynamic range in liquid media with little-to-no detectable activity in the absence of inducer. To achieve this, our system design includes two *tetO* operator sites to regulate the inducible *P_tet_* promoter, reducing leaky expression without compromising a high dynamic range. In order to provide consistent regulation and reduce heterogeneity, we created a codon-optimized *tetR(B)* repressor for *B. subtilis* and placed it under an organism-specific promoter for constitutive expression. The resulting system was cloned into a shuttle vector for stable single-copy chromosomal integration at the commonly used *amyE* locus. Together, these elements generated a system that balances tight repression and strong induction across diverse growth conditions.

Compared to existing systems, the refined promoter-repressor architecture we developed here offers improved sensitivity, enabling lower concentrations of tetracycline inducers to be used, including cTc and dox. Prior systems report using the inducer anhydrotetracycline (aTc) at starting concentrations ≥100-fold higher than those used here (*11, 51*). The enhanced sensitivity of our system enabled the use of alternative tetracycline family antibiotics with lower binding affinities to TetR (*52, 53*), and use of their specific dose-response characteristics. We found this was particularly advantageous for multicellular induction of colonies on agar plates, possibly due to the higher concentration of inducer used. This new capability may allow for expanded applications such as screening and selection of genetic libraries using bacterial colonies with a non-metabolizable and non-toxic inducer.

By characterizing the induction system in response to these three tetracyclines, we demonstrated∼consistent performance in rich and minimal media as well as during colony growth on agar plates. And despite differences in potency and toxicity, flow cytometry showed unimodal induction across all inducers’ operative concentrations, indicating that each inducer results in ∼homogeneous gene expression. The differences in induction properties and non-toxic ranges highlight clear trade-offs between the tetracyclines, with variation in titration windows and toxicity dictating suitability for different applications. The commonly used aTc remains the best choice for achieving maximal, non-toxic expression, offering sensitive induction at low concentrations. However, its extremely low effective range correlated with less reliable induction on solid media, where diffusion limitations and uneven exposure may be particularly important (Fig. 3A). In this context, cTc and dox provide a solution, by allowing graded induction in colonies. While cTc and dox require higher concentrations than aTc for comparable induction and exhibit greater toxicity at their upper limits, their broader titration windows provide more graded control. This tunability may make them valuable for applications requiring fine-tuned incremental control, such as balancing metabolic fluxes or mitigating overexpression toxicity (*54–56*). Higher tetracycline concentrations also reduce impact of operational concerns such as pipetting variability and impact of inducer purity and degradation in extended experiments. Additionally, each tetracycline varies in stability: dox is relatively stable under long-term culture conditions, while both aTc and cTc are more prone to degradation under light (*27*), and cTc more sensitive to pH as well (*57, 58*). These functional and logistical trade-offs allow this induction system to be adapted to specific experimental or industrial needs.

Together, in this work, we present a high sensitivity multi-tetracycline inducible gene expression system for *B. subtilis*. Expanding the system to multi-inducer compatibility and measuring its performance across growth and media conditions suggest it can be used in both basic and applied contexts where new, quantitatively characterized, media ∼agnostic systems are needed. The system is well suited for integration into complex synthetic circuits requiring quantitative modeling of gene expression dynamics, and compatibility with high-throughput fluorescence-based methods, including flow cytometry and colony-based screening and selection assays. Its performance in colonies on solid media demonstrates its potential for adaptability when multicellular control is needed and where other nutrient-based induction systems can perturb physiology. In the future, the promoter design of this system can be adapted to other developmental states and transcriptional programs, such as biofilms and sporulation where the regulatory systems are currently much more limited (*59*). Overall, this induction system creates a new, tightly regulated and high dynamic range multi-tetracycline responsive induction system for *B. subtilis* with design features that allow for future application-specific development.

## MATERIALS & METHODS

### Strain construction

Strains used in this paper are detailed in Table S1. *B. subtilis* (*Bs*168 *trpC2*) derived strains were made competent through the previously described two-step method (*60*). For chromosomal integration, plasmids were transformed into *B. subtilis* competent cells using standard techniques (*60*), successful transformants selected for on LB lennox plates (Fisher LB agar 50-488-752, Casein Peptone 10 g/L, Sodium Chloride 5 g/L, Yeast Extract 5 g/L, Agar 12 g/L) containing 100 µg/mL spectinomycin at 37 °C. Integration at the *amyE* chromosomal locus was verified using a starch-iodine test.

### Plasmid construction & cloning

Plasmids, oligos and sequences used in this work can be found in Supplemental Tables S2-S4. The *E. coli tetR* gene (class B from transposon Tn10) was codon optimized for *B. subtilis* using Integrated DNA Technologies (IDT) online tool with additional design changes made by hand to reduce potential internal start codons and ribosomal binding sites (RBS) prior to ordering through Azenta.

All plasmids were cloned and propagated in *E. coli* (NEB DH5α, C2987H) and successful strains selected for on LB lennox agar (Fisher LB agar 50-488-752, Casein Peptone 10 g/L, Sodium Chloride 5 g/L, Yeast Extract 5 g/L, Agar 12 g/L) plates containing 100 µg/mL ampicillin (Sigma Aldrich A9518-26G). Plasmid sequences were confirmed using sanger sequencing (Azenta).

### Media, induction & culture conditions

Minimal glucose media (S7 media: 1X MOPS Minimal Media (Teknova M2106), 1.32 mM Dipotassium phosphate, 1% Glucose, 0.1% Glutamic acid, 40 μg/mL Tryptophan) and rich Casein Hydrolysate media (CH media: Casein hydrolysate 10 g/L, L-glutamic acid (monosodium) 3.68 g/L, L-alanine 1.25 g/L, L-asparagine 1.39 g/L, Potassium phosphate monobasic 1.36 g/L, Ammonium chloride 1.34 g/L, Sodium sulfate 0.11 g/L, Ammonium nitrate 0.1 g/L, Ferric chloride hexahydrate 1 mg/L, Magnesium sulfate 48 mg/L, Calcium chloride dihydrate 25.6 mg/L, Manganese(II) sulfate monohydrate 22 mg/L, Tryptophan 20 mg/L) were made fresh every use. Stock solutions of the tetracyclines were prepared as follows: Anhydrotetracycline (Fisher 13803- 65-1) 8 mg/mL in 100% ethanol, and chlortetracycline hydrochloride (ThermoFisher J60095.14) and doxycycline hyclate (ThermoFisher J60422.06) 5 mg/mL in 50% ethanol.

*B. subtilis* strains were streaked out from frozen stocks for single colonies non-selectively on LB lennox agar plates and grown overnight at 37 °C. Single colonies were then used to inoculate 2 mL of media as indicated. Liquid cultures were grown with aeration to ∼mid-log phase, OD_600_∼0.2-0.4 at 37 °C). Each tetracycline inducer (aTc, cTc, or dox) was serially diluted from the respective stock solution to the concentration indicated in the appropriate media.

### Plate reader measurements & data analysis

For microplate experiments, the mid-log cultures were diluted 1:30 in triplicate into media with or without tetracycline in a 96-well black-wall, clear flat bottom plate (Greiner 655090). A BioTek Synergy Neo2 plate reader was set for continuous orbital shaking (425 cpm, 3 mm) at 37 °C. OD_600_ and *mCherry* (excitation 560/20 nm, emission 620/15, using optical filters) fluorescence measurements were taken every 12 min for 16 hrs.

In each experiment background values from media-only wells were averaged and subtracted from sample containing wells. The contribution of cellular autofluorescence at mid-log was subtracted using the autofluorescence values from the nonfluorescent strain (*Bs*168 *trpC2*). Fluorescence at∼ mid-log phase was averaged for every concentration across replicates. The nonlinear regression curves for each tetracycline dose-response and corresponding EC50 values were obtained using a fit to the Michaelis Menten equation (GraphPad Prism). Doubling times were calculated using an exponential fit for OD_600_ values over time in mid-log phase.

### Flow-cytometry for minimal media induction profiles

*B. subtilis* strains were grown as described above. When the cultures in the 96-well plates reached approximate mid-log, the plate was transferred to a Beckman Coulter Cytoflex S Cell Analyzer for single-cell measurements. Measurements for ∼50,000 events were taken per well. mCherry fluorescence was detected using the ECD channel (excitation: 561 nm, emission: 610/20 nm).

### Petri-dish induction & imaging

Minimal glucose cultures were inoculated from single colonies and grown to OD_600_ ∼ 0.1. Then 10 µL was spotted onto minimal glucose agar plates (± tetracycline as indicated) and allowed to dry at room temperature before incubating overnight (∼16 hrs) at 37 °C. Fluorescence images were taken using a BioTek Cytation5 using the Texas Red filter cube (ex 586/15 nm, em 647/57 nm, DC 605nm) using an integration time of 150 ms and a gain of 24. (Fiji Is Just) ImageJ (*61*) was used to background subtract the mean intensity of the petri dish from every pixel value for each respective image. A reference brightness range was manually set and normalized across images.

To calculate fold-change, the mean pixel intensity was quantified from each colony using (Fiji Is Just) ImageJ. To account for autofluorescence, the mean intensity of the non-fluorescent strain *Bs*168 *trpC2* was subtracted at each tetracycline concentration.

**Abbreviations**
anhydrotetracycline: (aTc)
chlortetracycline: (cTc)
doxycycline: (dox)
half-maximal response effective concentration: (EC50)

## Author Contributions

KRV designed, performed experiments, and analyzed data. EAL conceived and designed the study, supervised the project, obtained funding and resources. KRV and EAL wrote and edited the manuscript.

## Conflict of Interest

The authors declare no competing interests.

## Acknowledgements

This work was supported by NIH grant R35GM147429

We thank the Institute for Chemical Imaging of Living Systems (RRID:SCR_022681) at Northeastern University for consultation and instrument support.

## Data Availability Statement

The data that support the findings of this study are available from the author upon reasonable request. Genetic constructs will be deposited to a stock center upon publication.

